# Neural correlates of place learning in the cane toad, *Rhinella marina*

**DOI:** 10.64898/2026.08.31.748259

**Authors:** Daniel A. Shaykevich, Shuyun A. Xiao, Chloe L. Golde, Daniel W. Sorenson, Lauren A. O’Connell

## Abstract

Learning locations by integrating external reference points, or place learning, is a form of navigation strongly associated with the mammalian hippocampus and its homologs in other vertebrates. In contrast, cue learning relies on individual environmental features and can occur independently of the hippocampus. Despite growing interest in amphibian spatial cognition and association of spatial functions with the medial pallium brain region, how amphibian brains support different spatial learning strategies remains unknown. We designed a four-arm maze to test place, cue, and turn-direction learning in the cane toad (*Rhinella marina*), and paired maze trials with visualization of brain activity through pS6 immunohistochemistry. Toads improved performance over time only in the place task but exhibited consistently faster exit times in the cue task. Place task toads displayed elevated brain activity only in the dorsal and lateral palliums. These observations demonstrate that amphibians can acquire spatial navigation strategies through distinct behavioral paradigms and reveal the diverse pallial contributions underlying vertebrate spatial cognition.

## Introduction

Spatial memory is a vital cognitive function that supports behaviors critical to survival and fitness across animals (1–3). Successful navigation requires animals to acquire and use spatial information from their environment through multiple behavioral strategies. Some strategies rely on integrating relationships among environmental landmarks to determine a goal location within an allocentric framework (*place learning*) (4–6), whereas others depend on individual visual features (*cue learning*) (4,7,8) or stereotyped motor responses, such as maintaining a consistent turn direction (9,10). These navigation strategies differ in their cognitive demands and rely on distinct neural mechanisms (11,12).

The mammalian hippocampus is the best-studied vertebrate brain region involved in encoding spatial information (13,14), and homologous brain regions have been identified across vertebrates (11,15–17). In mammals, hippocampal regions communicate with brain structures with complementary roles, such as the entorhinal cortex (18). In some vertebrates, specific navigation tasks occur independently of the hippocampal homolog (11,12,19), especially cue-based tasks involving beaconing to a visual feature. However, how different navigation strategies are acquired and supported by neural circuits outside mammals remains poorly understood. Despite hints from comparative neuroanatomy and growing research in more diverse models, little is known about which brain regions support hippocampus-independent spatial tasks in non-mammalian vertebrates.

The neural basis of amphibian spatial cognition is in the early stages of characterization. Laboratory studies (16,20–22) and investigations on homing and orientation in the wild (17,23) have identified several regions associated with navigation, including the medial pallium– the amphibian hippocampal homolog– as well as the septum and lateral and dorsal palliums (16,17). While laboratory work has explored how the brain supports navigation related to environmental geometry and acoustic cues (20,21), the neural ensembles underlying visually guided place and cue learning in amphibians remain unresolved. As amphibians mark a critical transition in vertebrate evolution from aquatic to terrestrial environments, their spatial cognition remains a significant gap in the understanding of vertebrate neural evolution.

The cane toad (*Rhinella marina*) is an emerging model for amphibian spatial cognition (17,24,25). Although long distance navigation and homing behavior have been characterized in cane toads (17,25), the types of information they use to navigate at local spatial scales and the neural correlates supporting these behaviors remain unclear. To address these questions, we designed a four-arm maze (Supplemental figure S1) to test whether the cane toad can find a goal location through place learning, cue learning, or turn direction. Following maze training, we sampled brain tissue and quantified neural activity in telencephalic structures using pS6 immunohistochemistry. By comparing behavioral performance and regional patterns of neural activity across navigation strategies, this study provides a framework for understanding how different forms of spatial information support local navigation and are represented in the amphibian pallium.

## Materials and Methods

### Animals

Adult toads (n = 35, age unknown) were collected in West Palm Beach, Florida and housed at Stanford University. Toads were group-housed in glass terraria maintained at 20-25 and 80-100% humidity on a reverse light cycle to facilitate experiments. Toads were fed crickets three times weekly with ad libitum access to water. Two days before acclimatization trials, animals were housed individually in terraria. From this point, toads were no longer fed or provided water. Toad mass was monitored daily to ensure that it did not drop below 80% of pre-dehydration period mass. Toads weighed between 67 and 182 grams. Attempts were made to distribute toad sizes equally across groups, but cue toads were larger than place toads (Kruskal-Wallis: χ²_(3)_ = 10.91, p = 0.01, Supplemental figure S2). Both male and female toads were used but sex differences were not analyzed as previous experiments (17,20) and maze prototypes did not find differences in spatial behavior and/or brain activity.

### Maze Design

The four arm maze (Supplemental figure S1) was constructed out of acrylic (TAP Plastics, San Jose, CA, USA) and cut to size in a laser cutter (Glowforge, Seattle, WA, USA). The floor was made from opaque, 1/8” white acrylic and the walls and ceiling were made from clear, 1/8” acrylic. Each arm was 18” long, 5” wide, and 5” tall. Cut pieces were bonded with acrylic cement.

The maze was modelled on Rodríguez et al., 2002 (11) and adjusted for size and design based on preliminary experiments. Four identical arms were arranged at right angles leading into a central 5 in^3^ hub. In each trial, one arm was blocked, while the arm opposite was the entry arm. The remaining two were the potential exits. For each trial, depending on the toad’s training condition, one exit arm was “correct” and one was “incorrect”. Individual arms could be switched into any position to prevent bias for specific arms.

The maze was set in the middle of an experimental room on artificial grass. A heating pad was placed under the maze to warm the floor and encourage movement. The room was not uniform, so the visual environment around the maze differed and tape markings were put on the walls to increase variation. The short ends of the room were covered by curtains where experimenters could sit, and a GoPro camera was mounted on a rod over the maze. In each corner of the room, there was a Pyrex dish filled with water the animal could reach once it exited. An additional 5” cube made from opaque, white acrylic with one open side held toads between trials and introduced them into the maze.

### Training Conditions

Animals were trained in four conditions (Figure 1A). The entry side was randomly selected for each trial. “Place” condition animals (n = 9) were trained to locate a specific location: no matter what side of the maze they entered, the exit was a fixed location relative to external, allocentric landmarks, requiring either a left- or right-turn depending on the entry side for the trial. “Cue” conditions toads (n = 12) were trained to travel to a visual cue: an “X” on the door of the correct exit, which would randomly change positions between the two possible exits, so toads would have to turn either left or right depending from which side they were introduced. “Turn Direction” (TD) toads (n = 7) were trained to turn in the same direction for each trial (either left or right) irrespective of which side they entered the maze from. Lastly, the “Control” group (n = 7) were subject to trials where the exit was randomly selected and there was no visual identifier, so no fixed information allowed for learning.

**Figure 1.**
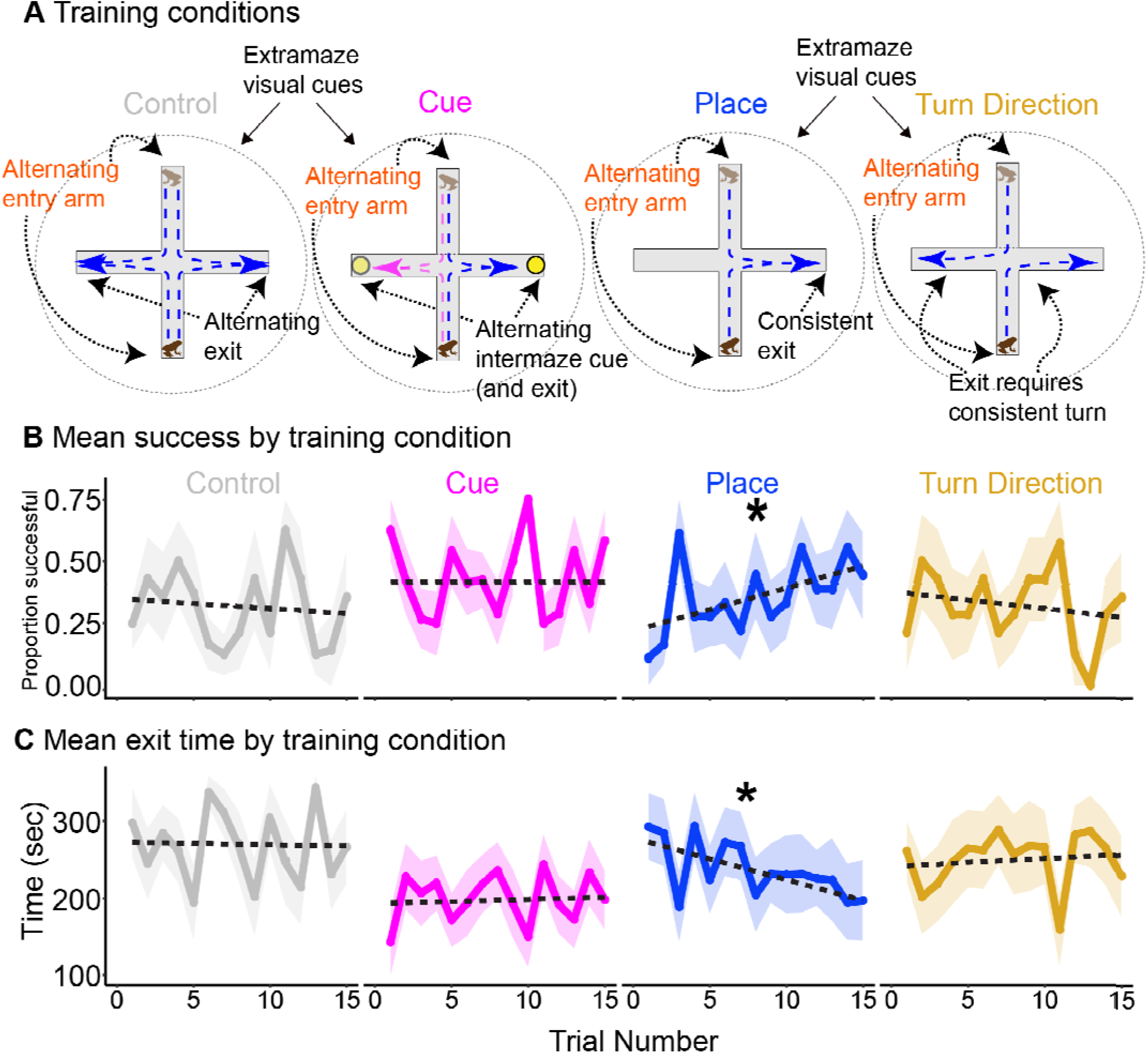
Toads in the place condition improved performance in the maze. **(A) Explanations of training conditions.** Control frogs had no consistent relationship between entry and exit arms. Cue toads were trained to exit through the arm marked by an intra-maze visual cue. Place toads could exit only through an arm that maintained a consistent spatial position. Turn Direction toads had to turn in a trained direction (either left or right) to exit. **(B) Learning curves depicting trial success.** Only place task toads showed a significant increase in successful trials over the 15 trial period (β = 0.0222 ± 0.0072 SE, t = 3.058, p = 0.0027). Shading represents standard error of the mean. **(C) Learning curves depicting maze exit times.** Only place task toads showed a significant decrease in the time it took to exit the maze over 15 trials (β = -5.506 ± 1.907 SE, t = -2.887, p = 0.0046). Shading represents standard error of the mean.

### Training Regimen

For two days prior to the beginning of training, animals underwent acclimatization trials, which were carried out by placing the toad into the maze with both doors open, and allowing it to move for five minutes. At the end of the trial, toads were guided to a water dish outside of the maze and allowed to soak for one minute.

Following two days of acclimation, toads performed five days of training trials, with three consecutive trials per day recorded with a GoPro camera. Trials took place from 12:00 – 15:00, at the end of the light cycle. At the beginning of each trial, the toad was placed into the opaque box constructed from white acrylic. The toad was released into the maze by hand and the observer retreated behind a curtain. The toad was allowed five minutes in the maze. If it did not exit, the toad was guided out of the correct exit by the experimenter and placed into the white box for two minutes before the next trial. If the toad exited, it was allowed to reach a water dish before being collected. In between animals, the entire maze was wiped down with 70% ethanol and allowed to dry.

### Behavior Coding

Maze performance was scored manually from recorded videos. Trials were scored with respect to escape time (time of 360 seconds assigned if toad did not leave within five minutes), direction of first turn (1 for right and 0 for left) and overall success (1 if the toad made the correct turn and left the maze, 0.5 if made the correct turn but did not leave immediately, and 0 if it did not make the correct turn).

### Tissue Collection

Thirty minutes after the completion of the last trial, animals were euthanized through subcutaneous injection with MS-222 (0.7% solution, 300 mg/kg body weight), and transcardically perfused with PBS followed by 4% paraformaldehyde (PFA). Brains were removed into 1 mL of 4% PFA and stored overnight at 4. The brain was washed three times in PBS (at least five minutes per wash at 4) and placed into 1 mL of 30% sucrose in PBS for cryoprotection. Once the brain sank in the sucrose solution, it was embedded in Tissue-Tek OCT Compound (Electron Microscopy Sciences) and frozen in a cryomold on dry ice before being stored at -80.

### Immunohistochemistry

Visualization of brain activity is identical to that in Shaykevich et. al 2025 (17). Brains were cryosectioned into 20 µm slices and thaw-mounted onto slides in four series. After drying at room temperature for ∼48 hours and storage at -80, we performed pS6 immunohistochemistry (Phospho-S6 (Ser244, Ser247) Polyclonal Antibody, Invitrogen, Waltham, MA, USA) to label translationally active neurons (26). We used a biotinylated secondary antibody (Goat Anti-Rat IgG Antibody (H+L), Vector Laboratories) and additionally performed cresyl violet staining for Nissl bodies in neurons. Slides were mounted with Fisher Chemical Permount Mounting Medium (ThermoFisher Scientific, Waltham, MA, USA).

### Microscopy and Cell Counting

Microscopy and cell counting were also similar to Shaykevich et al. 2025 (17). Slides were imaged on a Leica DM6B microscope with brightfield capability at 20x magnification. Three sections in the medial portion of the telencephalon were selected and used to count pS6-positive cells in each of six brain regions: medial pallium (Mp), lateral pallium (Lp), dorsal pallium (Dp), medial septum (Ms), lateral septum (Ls) and the striatum (Str). Both hemispheres of each section were imaged and counted. Delineation of brain regions was based on Sotelo et al. 2016 (20), which defined pallial regions by Westhoff and Roth 2002 (27) and ventral regions by Moreno and González 2004 (28). ImageJ was used to calculate the surface area of each region. Densities of active neurons were calculated by dividing the number of pS6-positive cells by the area.

### Data Analysis

All statistical analyses were performed in R Studio (v 2024.12.0 Build 467, Posit Software, PBC, Sunnyvale, CA) running R (v 4.4.2, R Foundation for Statistical Computing, Vienna, Austria). To test if time to exit the maze or success changed over trials within training conditions, we performed linear mixed effects models for each condition using the “lme4” package (v. 1.1-37) (29), where time in seconds or success score was the response variable, trial number was a fixed effect, and toad identity was a random effect. Additionally, to test if maze performance was related to training condition, we performed linear mixed models where time in seconds or success score was the response variable, and trial number, condition, and their interactions were fixed effects, and toad identity was a random effect.

We analyzed cell count activity as described in Shaykevich et al. 2025 (17). Briefly, we generated generalized linear mixed models (‘glmmTMB’ v. 1.1.11 in R) (30) to test for differences in brain activity and its relation with spatial learning conditions, using training condition, brain region and their interactions as the main predictors. Toad identity was included as a random effect to account for multiple sections considered per individual and brain region area was included as an offset variable to account for brain area size. Post hoc pairwise contrasts between training were calculated with estimated marginal means using the ‘emmeans’ package in R (v. 1.11.1) (31). We only considered brains of toads that performed above chance levels on the last day of maze trials, which constricted sample size (place: n = 6; cue: n = 5). No turn direction toads achieved the criteria and were not included in brain activity analysis. All seven control toads were included.

## Results

### Place trained animals improve performance over time

We tested if toads could learn a goal location using allocentric landmarks (place), a single visual feature (cue), or a consistent turn direction (Figure 1A). We first tested whether changes in trial success over the course of training differed among training conditions. Toads in the place condition displayed a greater increase in success over time than other groups (Supplemental table S1; Trial × Place interaction: β = 0.153±0.069 SE, z = 2.22, p = 0.027). Place toads exhibited the greatest decrease in maze exit time across training, although this relationship was not statistically significant (Supplemental table S2; Trial × Place interaction: β = −5.23±3.24 SE, t = −1.62, p = 0.107).

We next asked which conditions showed evidence of learning over training. Toads in the place condition showed increased trial success (Figure 1B; β = 0.0222 ± 0.0072 SE, t = 3.058, p = 0.003) and decreased exit time (Figure 1C; β = -5.506 ± 1.907 SE, t = -2.8887, p= 0.005) over successive trials (Supplemental table S3). No other group showed a significant change in success or exit time over the experiment. However, toads in the cue learning condition exhibited faster exit times across training than toads in control and turn direction conditions (Kruskal-Wallis: χ²_(3)_ = 18.037, p = 0.0004, Supplemental figure S3). Turning direction trained toads did not exhibit signs of learning, although all groups showed a bias for right hand turns (Supplemental figure S4).

### Pallial activity varies with training condition

We tested whether regional telencephalic brain activity differed among experimental conditions. All turn direction trained toads were excluded from analysis of brain activity because none met learning criteria on the last day. We found that training condition affected neural activity in a region-specific manner (Figure 2; brain region × training condition: χ²_(10)_ = 44.10, *p* = 3.15e−06). Specifically, toads in the place learning condition exhibited greater neural activity in the dorsal pallium (β = -0.90, SE = 0.31, z = -2.910, p = 0.01) and lateral pallium (β = -0.77, SE = 0.31, z = -2.52, p = 0.031) than toads in the control group (Figure 2B-C, Supplementary table S4). No other brain regions were significantly different across conditions.

**Figure 2.**
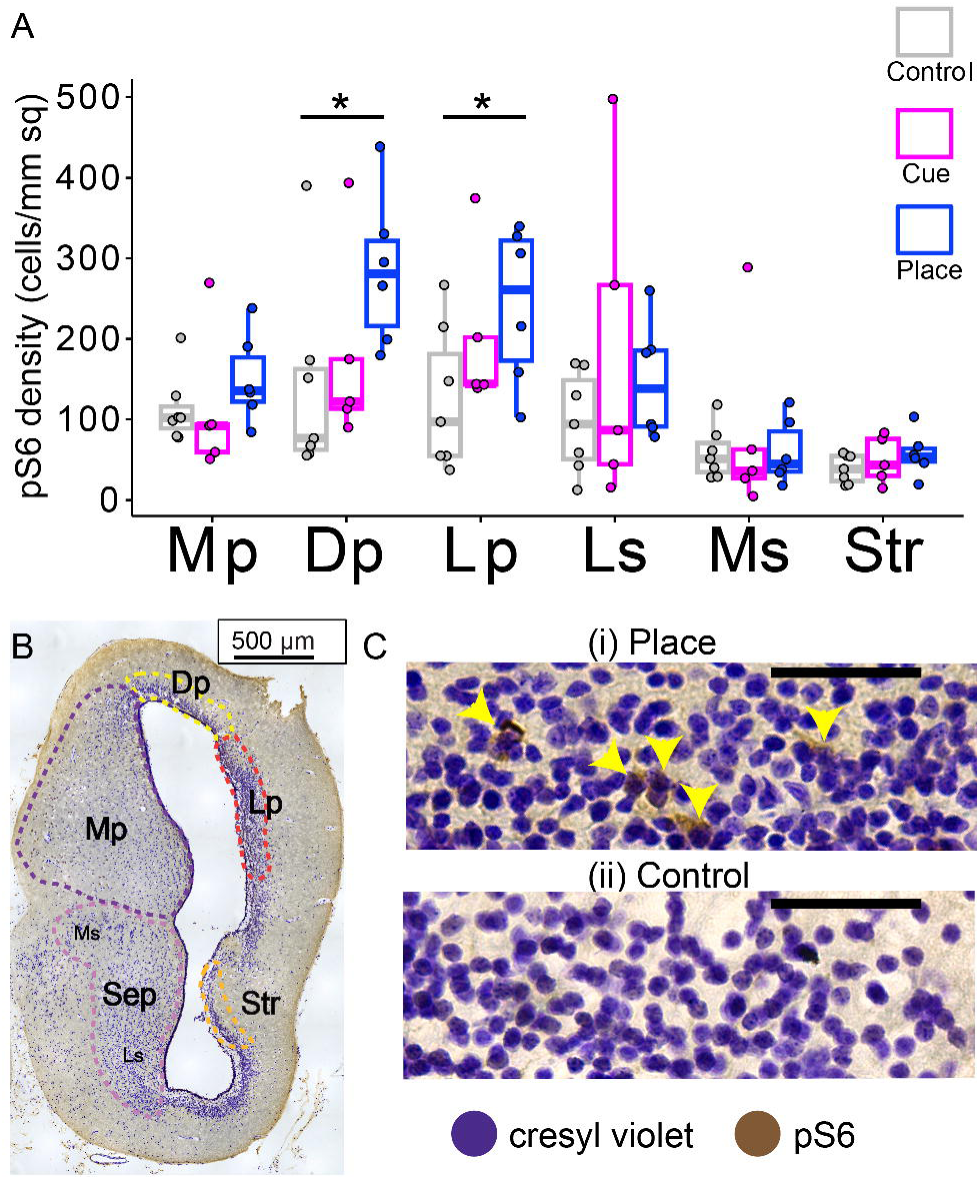
Place condition toads exhibit elevated activity in dorsal and lateral palliums. **(A) pS6-positive cells per mm^2^ averaged per individual for each brain region across training conditions.** Regional brain activity varied significantly due to training conditions (χ²_(10)_ = 44.10, *p* = 3.15e−06). Stars represent significant pairwise differences within brain regions based on a mixed model with cell count as a response factor. Abbreviations: Dp, dorsal pallium; Lp, lateral pallium; Mp, medial pallium; Sep, septum; Ms, medial septum; Ls, lateral septum; Str, striatum. (**B) Single hemisphere of representative toad brain slice with brain regions demarcated**. **(C) Representative examples of stained Dp brain tissue** from (i) place and (ii) control condition toads. Arrows indicate pS6-positive cells (brown stain). Cresyl violet stain is purple. Scale bars represent 50 µm.

## Discussion

Our findings demonstrate that toads can improve maze performance when trained using allocentric information but exhibit some potential taxis to visual cues that enable them to find exits faster. Differences in behavioral performance are accompanied by region-specific differences in pallial neural activity, but interestingly not by differences in activity within the medial pallium, the amphibian homolog of the mammalian hippocampus.

Toads in the place task showed incremental improvements in performance across training, but no evidence of gradual improvement was observed in any other training conditions. This finding extends previous evidence of spatial cognition in amphibians, including a poison frog (*Dendorabates auratus*) that was able to navigate a modified Morris-Water-Maze based on visual landmarks (32). The results suggest that, in addition to using distal or gradient based cues (like magnetoreception or olfaction) to travel long distances (33), amphibians can orient locally using proximal landmarks, in addition to other visual information like environmental geometry (20). Given the generally small scale of their natural movements (34) and repeated use of the same foraging grounds (35), cane toads may rely on local visual information for navigation at smaller spatial scales. Accumulating evidence suggests that amphibians can use diverse sensory information to navigate across different scales, similar to models suggested in other vertebrates (36).

Even though success rates and maze exit time did not change over the experiment, cue task toads exhibited faster exit times throughout training compared with toads in other conditions. In addition, on the final day of experiments, both place and cue condition toads showed higher levels of success than control and turn-direction toads (supplemental figure S5). These observations suggest that the presence of a distinct visual cue can immediately facilitate successful navigation, potentially through spontaneous orientation toward salient visual features. Naive taxis towards visual stimuli has also been observed in fish that display spontaneous preference for certain landmarks and patterns (37,38). Future tests incorporating variable cues may determine if specific shapes, patterns, or movements associated with naturally occurring stimuli are attractive (or repellant) to toads (39–41).

Although turn direction toads did not show evidence of learning, toads across conditions displayed a bias toward making rightward turns. This phenomenon was also observed in maze prototypes and provides additional evidence for behavioral lateralization in toads (42). Lateralization (43) and turning bias (44) are well documented in amphibians and may result in development-related morphological or neural asymmetry (45). Thus, poor performance in this learning condition may have been influenced by innate lateralization. In addition, consistent turn direction is unlikely to represent a reliable navigation strategy in natural environments.

Contrary to our expectations, we did not observe differences in medial pallium activity across conditions. In other vertebrates, hippocampal function specifically supports place, but not cue, learning (11,12,19). However, a lesion study in the toad *R. arenarum* showed that, while the medial pallium is necessary for navigation based on environmental geometry, ablation of the region also disrupted orientation using a visual feature cue (22). Thus, our findings suggest that both place and cue learning may rely on the medial pallium and add to growing evidence that the region governs a wider range of spatially-related cognitive processes and does not correspond directly with the function of the mammalian hippocampus. This can potentially be attributed to different neural architecture, including more unprocessed sensory inputs than found in other vertebrate hippocampal regions (16,22,27,46).

We observed significant differences in the dorsal and lateral palliums, which are analogous to the mammalian allocortical regions and entorhinal and piriform cortices, respectively (16). The lateral pallium had higher activity in *R. marina* performing long distance homing (17) and both dorsal and lateral palliums had increased activity in *R. arenarum* navigating a geometry-based arena with a visual feature (20). Additionally, both regions showed elevated activity in wild poison frogs during orientation following displacement (23). Interestingly, we did not observe differences in activity in these regions in the cue condition; rather, we observed differences specifically between the control and place conditions. These observations suggest that dorsal and lateral pallial activity may be broadly involved in governing spatial tasks, rather than exclusively reflecting the use of a particular navigational strategy. Functional tests of spatial coding through electrophysiology would be required to fully delineate the specific contributions of the medial, dorsal, and lateral palliums with respect to amphibian navigation.

## Conclusion

Our study demonstrates that cane toads can learn to navigate locally through place learning, adding to evidence that amphibians are capable of determining their position relative to their surroundings across both small and large spatial scales. The dorsal and lateral palliums showed increased activity in the place condition compared to the control, suggesting that these regions are associated with spatial learning in amphibians. Together with previous experiments, these results suggest that homologous pallial regions may not exhibit direct functional equivalence across vertebrate lineages.

## Acknowledgements

We thank Erez Krimsy for his help in designing early versions of the maze. We also thank Nicholas Panyanouvong for performing preliminary trials that informed the final version of the experiment.

## Ethics

All procedures were approved by the Institutional Animal Care and Use Committee of Stanford University (Protocol #33530). Cane toads were housed at Stanford University under Restricted Species Permit No. 2352 from the California Natural Resources Agency, Department of Fish and Wildlife.

## Funding

This work was supported by the National Science Foundation CAREER award (IOS-1845651) and the National Institute of Health BRAIN Initiative (R34NS127103) to L.A.O. D.A.S. was supported by a National Science Foundation Graduate Research Fellowship (2019255752) and the National Institutes of Health (T32GM007276).

## Use of Artificial Intelligence

No AI tools were used in any aspect of this project, including writing or analysis.

## Data, code and materials

Data and R script have been uploaded with restricted access to Zenodo and will be made publicly available on publication: https://doi.org/10.5281/zenodo.22178372

## Competing interests

The authors have no competing interests.

## Credit

Conceptualization: DAS, LAO

Data curation: DAS

Formal analysis: DAS

Funding acquisition: LAO

Investigation: DAS, SAX, CG, DWS

Methodology: DAS

Project administration: LAO

Resources: LAO

Supervision: LAO

Visualization: DAS

Writing – original draft: DAS, LAO

Writing – review and editing: SAX, CG, DWS

## Notes

### Competing Interest Statement

The authors have declared no competing interest.

