## Supplemental Figures and Tables for "Neural correlates of place learning in the cane toad, *Rhinella marina*"

### Supplementary Information

Daniel A. Shaykevich[1‡*], Shuyun A. Xiao[1], Chloe L. Golde[1], Daniel W. Sorenson[1,2], and Lauren A. O'Connell[1*]

[1]Department of Biology, Stanford University, Stanford, CA, USA
[2]Department of Biological Sciences, San José State University, San Jose, CA, USA

[‡]Current Address: Department of Interdisciplinary Life Sciences, University of Veterinary Medicine, Vienna, Vienna, Austria

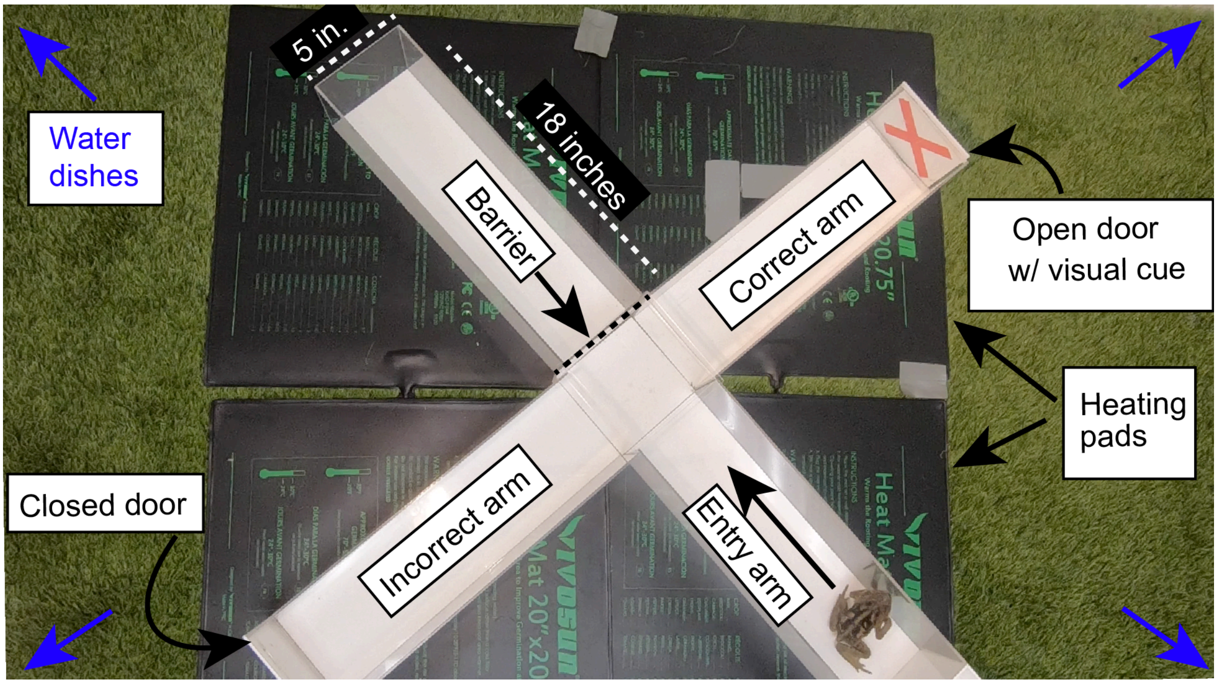


**Figure S1. Four arm maze design.** The maze consisted of four 18 inch by 5 inch arms that came together in a central hub. The maze sat on top of heating pads to warm the floor and under a bright light to increase toad motivation to move. The opposite side from the entry arm was blocked with a plastic barrier. The incorrect door was fixed to the maze with velcro while the correct door could be knocked down by the toad to exit. Pictured is the "X" used in the cue condition. Water dishes were located in the corner of the testing space so toads could reach them after exiting.

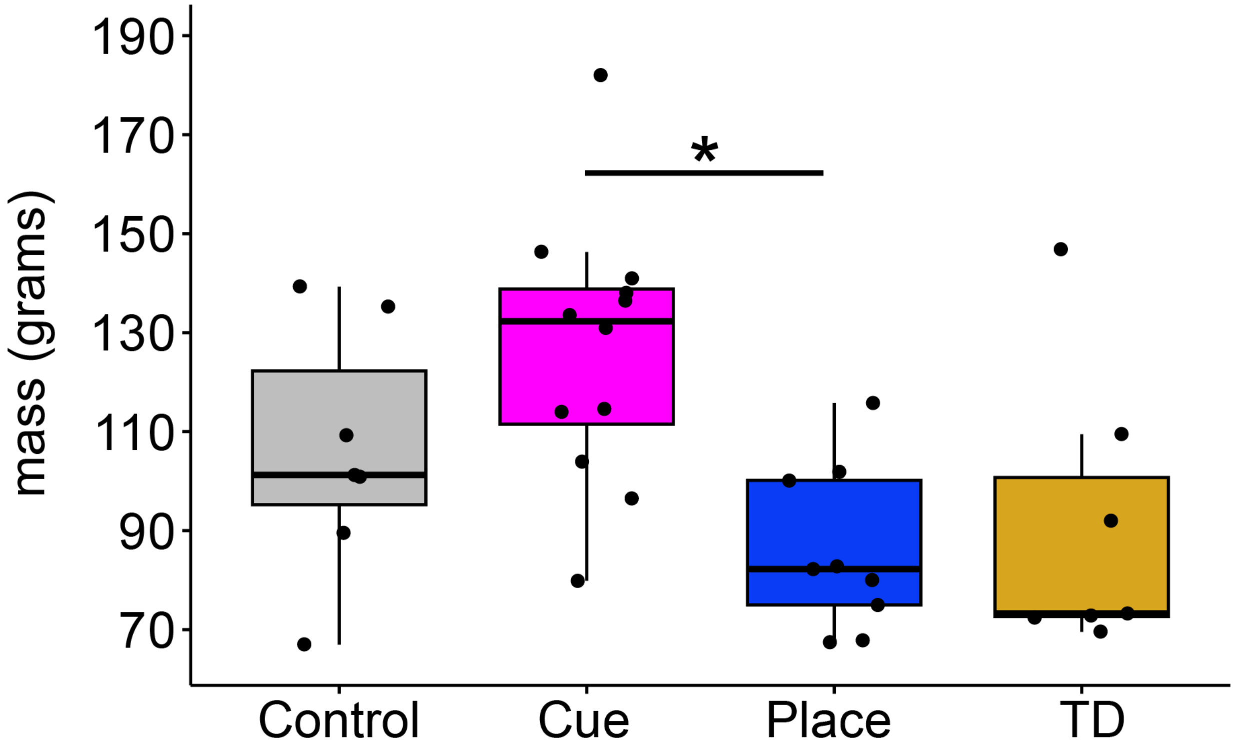


**Figure S2. Toad size differed between conditions.** Though toads were randomly selected for study, toads in the Cue condition were larger than toads in the Place condition (Kruskal-Wallis: $\chi^2_{(3)} = 10.91$, p = 0.01). Asterisks represent significant differences from post hoc Dunn's test (with Holm adjustment).

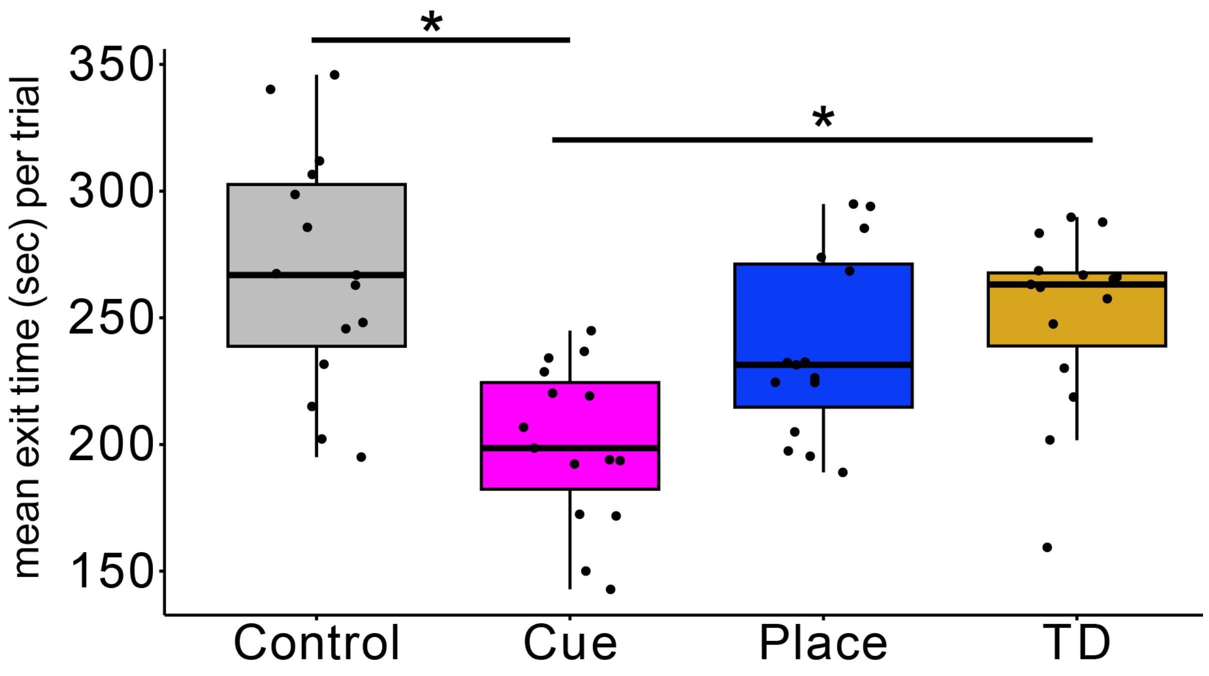


**Figure S3. Toads trained in the cue task generally left the maze faster.** Independent of trial success, these toads tended to find the maze exit and leave faster than other groups (Kruskal-Wallis: $\chi^2_{(3)} = 18.037$, p = 0.0004). Asterisks represent significant differences from post hoc Dunn's test (with Holm adjustment).

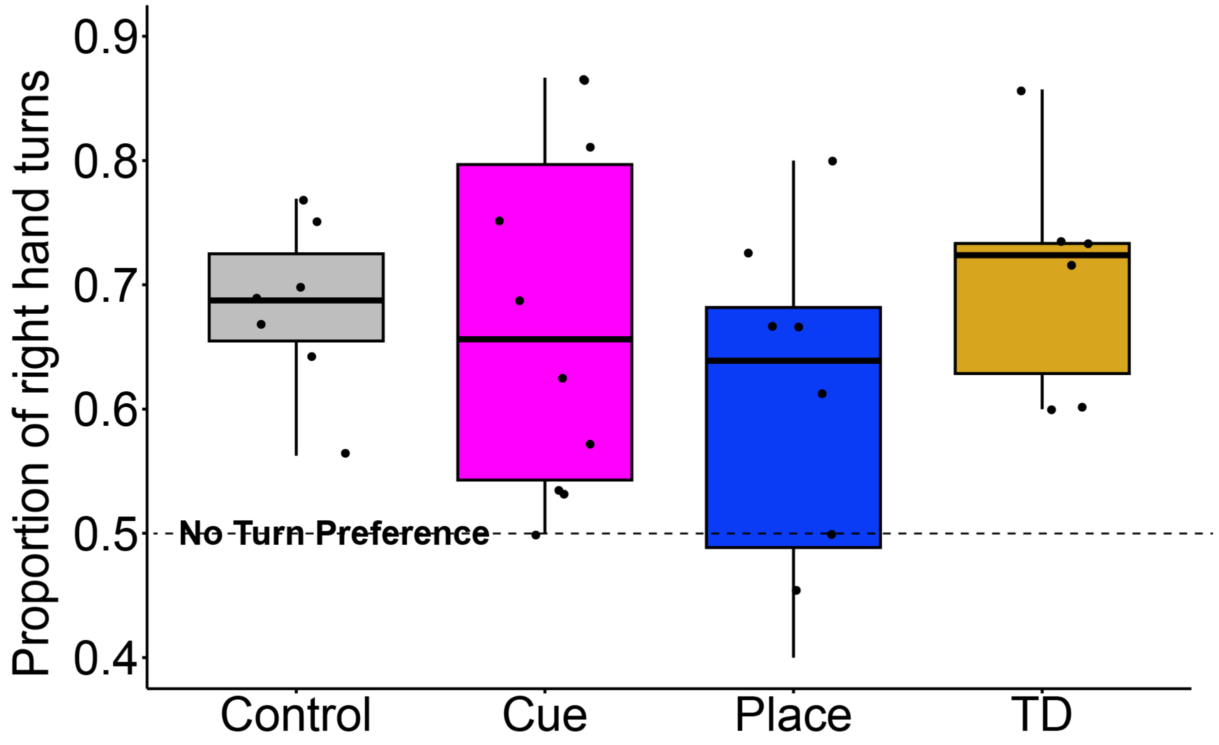


**Figure S4. Toads across all conditions show a right-turn bias.**. Each point represents the proportion of right hand turns made by each individual over 15 trials- turn preference of 0.5 would indicate equal amounts of left- and right- hand turns.

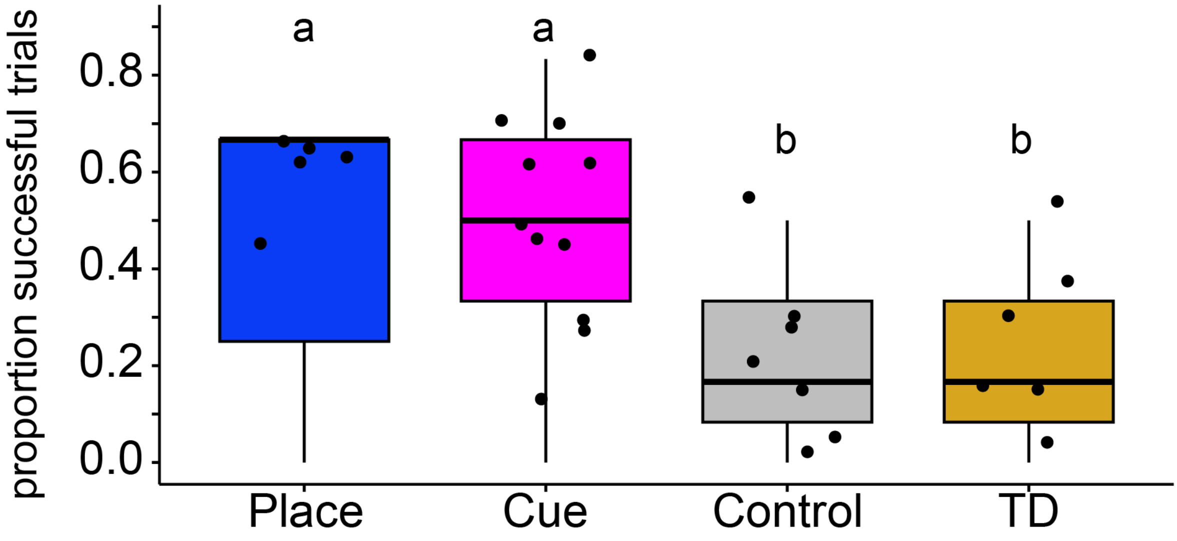


**Figure S5. On the last experimental day, place and cue toads showed higher levels of success.** Each point represents the proportion of successful trials on the last day of the experiment, with the place and cue toads outperforming control and turn direction toads (Kruskal-Wallis: $\chi^2_{(3)} = 9.96$, $p = 0.019$). Letters represent significance levels from post-hoc Dunn's test using unadjusted p-values. P-values adjusted with the Benjamin-Hochberg method show marginal differences between control and both place and cue toads p = 0.051 and p = 0.591, respectively), and cue and turn direction toads (p = 0.079).

**Table S1. Mixed-effects linear modeling success as a function of trial number**

| Fixed Effects | Estimate | Std. Error | z value | p-value |
|---|---|---|---|---|
| Trial | -0.02268 | 0.05323 | -0.426 | 0.6701 |
| ConditionCue | 0.42343 | 0.57640 | 0.735 | 0.4626 |
| ConditionPlace | -1.06082 | 0.66067 | -1.606 | 0.1083 |
| ConditionTD | 0.25057 | 0.66348 | 0.378 | 0.7057 |
| Trial:ConditionCue | 0.02074 | 0.06414 | 0.323 | 0.7464 |
| Trial:ConditionPlace | 0.15258 | 0.06884 | 2.217 | **0.0267** * |
| Trial:ConditionTD | -0.03515 | 0.07564 | -0.465 | 0.6421 |

***Significant relationships**

**Table S2. Mixed-effects linear modeling exit time as a function of trial number**

| Fixed Effects | Estimate | Std. Error | df | t value | p-value |
|---|---|---|---|---|---|
| (Intercept) | 270.1051 | 36.0156 | 64.5857 | 7.500 | 2.34e-10 |
| Trial | -0.2489 | 2.5029 | 501.3809 | -0.099 | 0.921 |
| ConditionCue | -74.1347 | 45.2311 | 64.1254 | -1.639 | 0.106 |
| ConditionPlace | 10.9173 | 47.7749 | 63.3227 | 0.229 | 0.820 |
| ConditionTD | -31.9664 | 50.8121 | 64.0188 | -0.629 | 0.532 |
| Trial:ConditionCue | 0.9217 | 3.1443 | 501.2521 | 0.293 | 0.770 |
| Trial:ConditionPlace | -5.2294 | 3.2375 | 502.1669 | -1.615 | 0.107 |
| Trial:ConditionTD | 1.8350 | 3.5252 | 501.2492 | 0.521 | 0.603 |

**Table S3. Mixed effects linear models measuring performance over trials within conditions**

| Condition | Variable | Fixed Effects | Estimate | Std. Error | df | t-value | p-value |
|---|---|---|---|---|---|---|---|
| Control | Time | (Intercept) | 270.0482 | 28.3261 | 22.6881 | 9.534 | 2.13e-09 |
| | | Trial | -0.2552 | 2.4756 | 101.3488 | -0.103 | 0.918 |
| Control | Success | (Intercept) | 0.366166 | 0.085493 | 107.0 | 4.283 | 4.04e-05 |
| | | Trial | -0.00560 | 0.009388 | 107.0 | -0.597 | 0.552 |
| Cue | Time | (Intercept) | 195.9853 | 28.9765 | 21.5897 | 6.764 | 9.39e-07 |
| | | Trial | 0.6716 | 1.9685 | 170.0159 | 0.341 | 0.733 |
| Cue | Success | (Intercept) | 0.4114 | 0.07375 | 85.25 | 5.578 | 2.82e-07 |
| | | Trial | 0.002464 | 0.007696 | 170.3 | 0.320 | 0.749 |
| Place | Time | (Intercept) | 281.183 | 35.739 | 12.322 | 7.868 | 3.77e-06 |
| | | Trial | -5.506 | 1.907 | 131.326 | -2.887 | ***0.00455** |
| Place | Success | (Intercept) | 0.1918 | 0.09131 | 24.01 | 2.101 | 0.0463 |
| | | Trial | 0.02215 | 0.007244 | 132.1 | 3.058 | ***0.0027** |
| Turn Dir. | Time | (Intercept) | 238.359 | 31.765 | 17.375 | 7.504 | 7.53e-07 |
| | | Trial | 1.561 | 2.585 | 99.066 | 0.604 | 0.547 |
| Turn Dir. | Success | (Intercept) | 0.388907 | 0.092046 | 36.0905 | 4.225 | 0.000155 |
| | | Trial | -0.00834 | 0.009087 | 99.2161 | -0.917 | 0.361199 |

***Significant relationships**

**Table S4. Pairwise contrasts between training condition of brain region specific neural activity in toads that performed above chance level on final experimental day**

| Brain region | contrast | estimate | SE | z.ratio | p.value |
|---|---|---|---|---|---|
| DP | Control - Cue | -0.3295 | 0.328 | -1.004 | 0.5744 |
| DP | Control - Place | -0.8970 | 0.308 | -2.910 | **0.0101*** |
| DP | Cue - Place | -0.5675 | 0.338 | -1.681 | 0.2126 |
| LP | Control - Cue | -0.6425 | 0.326 | -1.973 | 0.1189 |
| LP | Control - Place | -0.7744 | 0.307 | -2.523 | **0.0312*** |
| LP | Cue - Place | -0.1319 | 0.336 | -0.393 | 0.9184 |
| Ls | Control - Cue | -0.2038 | 0.324 | -0.629 | 0.8044 |
| Ls | Control - Place | -0.5193 | 0.305 | -1.700 | 0.2051 |
| Ls | Cue - Place | -0.3155 | 0.335 | -0.943 | 0.6133 |
| MP | Control - Cue | 0.1105 | 0.323 | 0.342 | 0.9374 |
| MP | Control - Place | -0.2602 | 0.305 | -0.854 | 0.6692 |
| MP | Cue - Place | -0.3707 | 0.333 | -1.112 | 0.5067 |
| Ms | Control - Cue | -0.1247 | 0.330 | -0.378 | 0.9243 |
| Ms | Control - Place | -0.0474 | 0.311 | -0.153 | 0.9873 |
| Ms | Cue - Place | 0.0773 | 0.341 | 0.227 | 0.9721 |
| Str | Control - Cue | -0.0865 | 0.352 | -0.246 | 0.9673 |
| Str | Control - Place | -0.3729 | 0.329 | -1.134 | 0.4933 |
| Str | Cue - Place | -0.2864 | 0.362 | -0.791 | 0.7083 |

***Significant relationships**
